# Relative roles of genetic variation and phenotypic plasticity in the invasion of monkeyflower *Erythranthre gutatta* in New Zealand

**DOI:** 10.1101/2022.06.06.495034

**Authors:** Michelle Williamson, Daniel Gerhard, Philip Hulme, Hazel Chapman

## Abstract

Evolutionary processes which increase the probability of an introduced plant species becoming invasive include high levels of genetic diversity and phenotypic plasticity. Naturalised in New Zealand, monkeyflower, *(Erythranthre gutatta),* a clonally spreading herb of waterways and seepage areas native to the Western USA, shows marked variation in a range of vegetative, reproductive and inflorescence traits. We used two common gardens differing in elevation to explore the relative contribution of genetic versus plastic variation within nine traits among 34 monkeyflower clones from across the New Zealand South Island. We looked for evidence of clinal variation across elevation gradients and for home site advantage. We found both high genetic diversity and trait plasticity explain the observed variation, although less evidence for adaptive plasticity. Most genetic variation was observed in the lowland garden (9m a.s.l.), where the overall trend was for above ground dry weight to be lower, and horizontal shoot length greater, than at the montane garden (560m a.s.l). We found no evidence of local adaptation to any of the measured environmental variables. However, we observed a pattern of higher biomass and higher plasticity at lower versus higher elevations and in clones originating from lower elevation sites.

## INTRODUCTION

Invasive alien plants often show considerable phenotypic variation in their introduced range, explained by local adaptation, phenotypic plasticity (whereby plants are able to adjust their phenotype according to the local environment (Pigliucci et al. 1995; Geng et al. 2007; Pitchancourt and Van Klinken 2012) or a combination of both (Moroney et al. 2013; Si et al. 2014). However, despite a plethora of studies ( Geng et al. 2016; Hiatt and Flory, 2020; Fakhr et al. 2022;), the relative contribution of genes and the environment to such variation remains poorly understood (Vanwallendael et al., 2018; Villellas et al. 2021; Yuan et al. 2022). While the success of clonally reproducing invasive species is often attributed to plasticity (Loomis and Fishman 2009; Riis et al. 2010; Keser et al. 2014), quantitative links between plasticity and fitness are rare (Liu et al. 2016; Bufford and Hulme 2021a). Plastic traits can be neutral or even mal-adaptive, a consequence of genetic correlation or trade-offs (van Kleunen and Fischer 2004) and so evidence of plasticity does not necessarily equate to increased fitness. Of course, adaptation can play an important role in invasion success, even in clonal plants (Geng et al. 2016).

Given that understanding why some species become invasive but others don’t is one of the main focal areas of invasion biology (Moloney et al. 2009) this facet of invasion success warrants further investigation (Ghalambor et al. 2007; Geng et al. 2016; Lee-Yaw et al. 2019; Enders et al. 2020). It is especially relevant for clonal ornamental plants given their under-representation in the literature and the predicted global increase in garden escapes with climate change (van Kleunen et al. 2018; Hulme 2020). Many ornamentals reproduce easily, often with a strong clonal component to reproduction and are predicted to be the major pool of future invasive species worldwide (van Kleunen et al. 2018).

Local adaptation allows for invasive success (Lee et al. 2002; Leimu and Fischer 2008; Colautti and Lau 2015; Molina-Montenegro et al. 2018; Lee-Yaw et al. 2019) by conferring genotypes with a ‘home site advantage’ (Kawecki and Ebert 2004; Maron et al. 2004; Allan and Pannell 2011; Bennington et al. 2012), whereby plants grown in their home environment are more fit than plants from elsewhere (but see Lee-Yaw et al. 2019). In contrast phenotypic plasticity allows the same genotype to thrive in a range of environments (Bossdorf et al. 2005; Richards et al. 2006; Hulme 2007; Moloney et al. 2009,; Bufford and Hulme 2021a; but see Palacio-Lopez and Gianoli 2011). As with local adaptation, plasticity may evolve post introduction (Lande 2015). Demonstrating the relative contribution of each evolutionary strategy is difficult for a number of reasons (Noble et al. 2019; Villellas et al. 2021) and findings have to date been equivocal (Bossdorf et al. 2005; Richards 2006; Hulme 2007; Bock et al. 2015).

One approach to untangle the different components (genetic versus plastic) of intra-specific phenotypic variation is to use common garden experiments, where clones from a number of environmentally different populations are grown together under as near -identical conditions as possible (Allan and Pennell 2009; Cheplick 2015; Villellas et al. 2021). If, under the same environmental conditions clones maintain their home site phenotypes, this variation is deemed genetic (Turesson 1922; Nunez-Farfan and Schlichting 2001). In contrast, if under the same conditions the clones from different locations exhibit similar phenotypes the variation is considered plastic. Comparing trait values within and among clones, analysis of variance allocates the proportion of variance within clones into genetic and plastic components. In invasion ecology common garden experiments have been used to compare provenances from native and introduced ranges in the introduced range (Bufford and Hulme 2021a; b; Colautti and Barrett 2009 and references within), alien provenances from just the introduced range (Kollmann and Baneulos 2004; Rapson and Willson 1992; Colautti and Barrett 2013; Monty and Mahy 2009) and provenances in both the native and introduced range (Maron et al. 2004; Flory et al. 2011; Shelby et al. 2016; Berend et al. 2019; Oduor et al. 2016 and references therein). However, often such experiments have been compromised by their design. For example, too few gardens or gardens with very similar environments (Moloney et al. 2009; Villelas et al. 2021). Moreover, well designed experiments have shown that contradictory results in different gardens are to be expected and help to untangle genetic versus environmental influences (Maron et al. 2004; Williams et al. 2008; Moloney et al. 2009; Florey 2011; Villelas et al. 2021).

We investigate the relative contribution of genotype versus plasticity to the widespread success in New Zealand of the introduced *Erythranre gutatta* (D.C.) G. L. Nelson, (section *Simiolus,* Phrymaceae), previously known as *Mimulus gutattus* D.C. Lowry et al. (2019), and commonly referred to as monkeyflower. Due to its wide range of life histories and environmental tolerance, and its propensity for whole genome duplication, *E. gutatta* has become a model species in evolutionary ecology and invasion biology (Wu et al. 2008; Friedman et al. 2015; Da Re et al. 2020). *E. gutatta* is a widespread, common and rapidly spreading clonally reproducing weed in New Zealand and is of considerable interest because it shows a marked niche-shift, occupying habitats in New Zealand which lie outside of the environmental conditions found in its native range (Da Re et al. 2020). One explanation for this lack of niche conservatism is that different genotypes (different source populations) respond differently to different environments. Alternatively, the niche-shift may reflect post-introduction evolution, either of plasticity or by local adaptation (Da Re et al. 2020). While New Zealand *E. gutatta* populations undoubtedly represent bottlenecked populations and, according to (Vallejo-Marin et al. (2021) historical, rather than recent introductions, they do include both native US and bridgehead UK populations, which suggests the potential for novel genetic combinations in New Zealand. How likely this is remains speculative given the primarily clonal nature of *E. gutatta* spread.

We undertook a set of common garden experiments to distinguish the extent to which the considerably phenotypic variation observed in *E. gutatta* in New Zealand reflects plasticity versus contemporary evolution leading to local adaptation. The selection pressure we considered most likely to impact plants from across our sampled sites was elevation, incorporating a steep cline in temperature and rainfall. We thus chose our two common gardens based on elevation; Ilam (hereafter referred to as the lowland site) and Cass (hereafter referred to as the montane site), at 9 m and 580 m respectively, to expose any GxE effect (different genotypes responding differently to the two environments). Given that our sites spanned the entire length of the South Island, we also looked for evidence of a latitudinal cline in biomass. Our specific hypotheses were that 1) we would detect clinal variation associated with elevation, so that clones from montane populations would be fittest in the montane garden and vice versa. 2) We would detect home site advantage, such that the Cass clone grown in the Cass garden would be the fittest clone in the montane garden and similarly, clones from close to the lowland Ilam garden would be the fittest clones in the lowland garden.

## Methods

### Study species

*Erythranthre gutatta* (D.C.) G. L. Nelson, is the most common species in a large *Mimulus* species complex (Ritland and Ritland 1989). Widespread across the Western US (Friedman et al. 2015; Demarche et al. 2016), *E. gutatta* has become naturalised across Europe and in New Zealand. First recorded in New Zealand in 1878 (Webb et al. 1988), Vallejo-Marin et al. (2021) found that New Zealand populations have been introduced multiple times both directly from native Alaska and by way of a bridgehead from the UK.

### Experimental design

We sampled a single clone from each of 34 distinct locations across the South Island of New Zealand. Locations were never less than 10 km apart and were chosen to be representative of environmental and altitudinal variation across the South Island and included seven geographically isolated regions (Fig 1).

**Fig 1.**
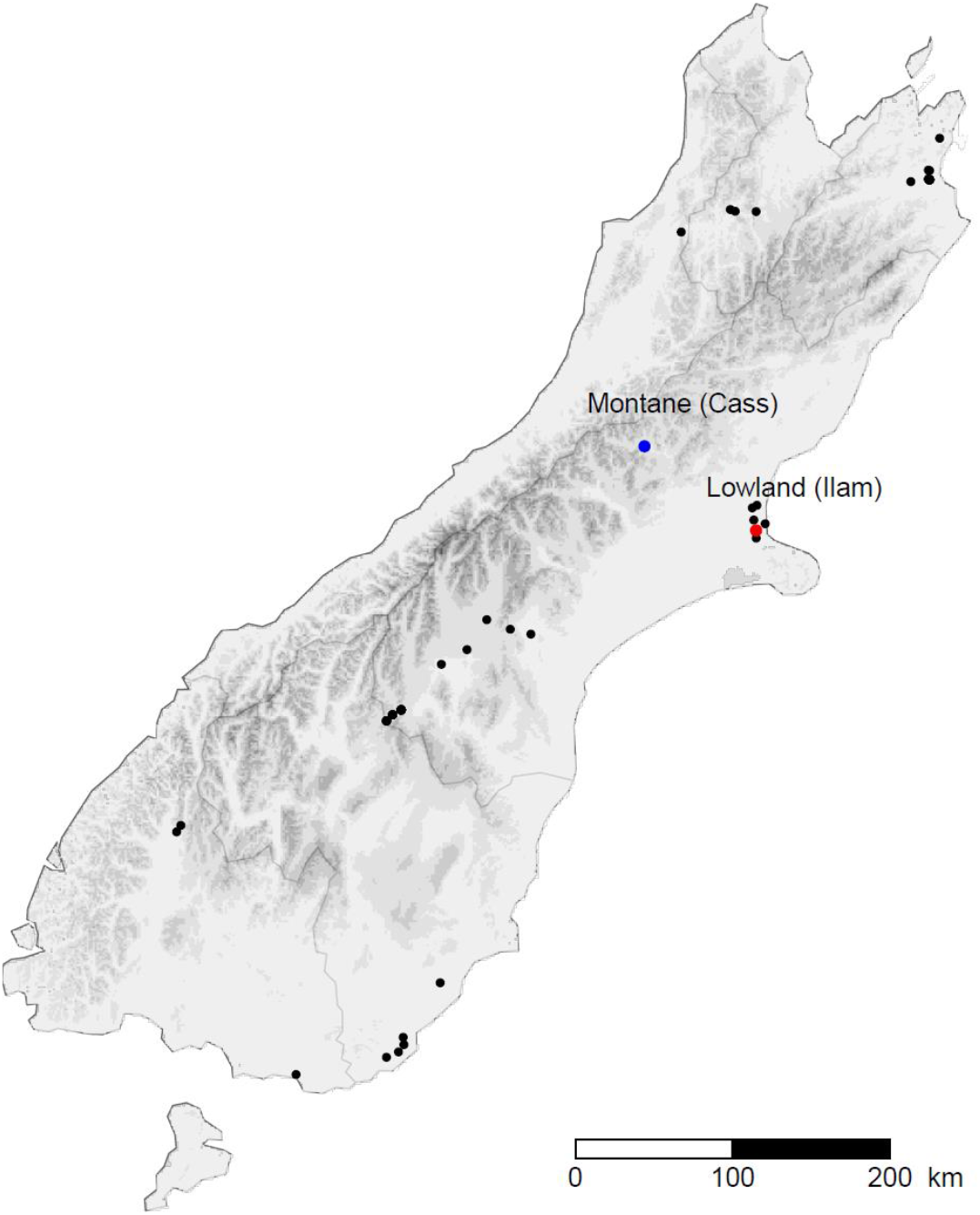
Sample locations of *E. gutatta* across the New Zealand South Island (https://www.stats.govt.nz). The sites are clustered according to geographical region.

At each sampling site we recorded 16 environmental variables and calculated the annual average temperature, growing season average temperature, average minimum and maximum temperature, extreme minimum and maximum temperature, annual average rainfall, humidity and precipitation from data collected from local climate stations between 1971 and 2000 (Table S 1). Populations were mostly small patches along stream sides or drainage ditches and we collected stems of *E. gutatta* with basal runners from a single clone from each location. From each clone we propagated twelve replicates from small internode cuttings. To reduce clonal environmental effects as much as possible, cuttings were <2cm long and were in similar developmental and physiological condition (Libby 1962; Weiner et al. 1997). Cuttings were planted in separate 10 × 10 × 10 cm^3^ pots filled with potting mix. Pots were placed on tables in a glasshouse, with their distribution on the tables randomized weekly. Early in the 2018 growing season cuttings were translocated into individual pots (7.5L, 20.5 cm wide x 25.5 cm diameter), lined with plastic bags and filled with a slow release fertiliser potting mix. The pots were moved into the two common gardens. Half (six replicates per clone) went into the lowland garden on the University of Canterbury Ilam campus (43°31’S 172°35’ and the remainder were set up at a montane garden located at the University of Canterbury Cass field station (43°02’S, 171°45’E). The montane garden was significantly cooler and wetter than the lowland garden (Table 1).

**Table 1.** Environmental measures of Ilam, Christchurch and the Cass Field Station. Averages 1971-2000

| Location | Elevation (m) | Annual Average Temperature (°C) | Average Temperature (°C) | Average Min Temperature(°C) | Extreme Min Temperature(°C) | Average Max Temperature(°C) | Extreme Max Temperature(°C) | Annual Average Rainfall (mm) | Humidity (%) | Precipitation (mm) | Max Sunlight Hours | Min Sunlight Hours |
| --- | --- | --- | --- | --- | --- | --- | --- | --- | --- | --- | --- | --- |
| Ilam, Christchurch | 9.69 | 12.29 | 14.7 | 9.63 | 0.62 | 19.76 | 32.5 | 131.3 | 75.3 | 10.10 | 15.25 | 8.27 |
| Cass Field Station | 578.81 | 8.22 | 10.72 | 4.17 | -5.3 | 17.27 | 30.06 | 2289.6 | 84 | 192.6 | 15.22 | 9 |

Plants were randomly assigned positions in each common garden and their distributions randomized weekly. E. *gutatta* occurs primarily in wetlands and riparian areas and thus plants were kept moist with daily automatic watering to limit any effect of drought or precipitation on performance.

### Performance traits

We chose nine quantitative traits which have previously been shown to exhibit marked inter-population variation in *E. gutatta* (eg Hall 2006; van Kleunen and Fischer 2008). We counted the maximum number of buds on each plant. In mid-April, towards the end of growing season, we measured the vegetative, reproductive and flower traits in both gardens: mean length and width of the largest two leaves, length of the longest horizontal shoot and length between the second and third internodes, the largest flower length, depth and width. All plants were then harvested towards the very end of the growing season in mid-April and dry weight of above ground tissue was measured after drying at 80°C for 72 h. We chose above ground dry weight as our proxy for fitness.

### Statistical analysis

To estimate the total genetic and environmental components of the variation in vegetative, reproductive and flower traits among clones, and their genotype x environment (GxE) interactions, we used linear mixed-effects models. In the models, each of the quantitative traits were analysed separately, using a logarithmic transformation on each observation after visually inspecting model diagnostics. For the GxE investigation for each response, the expected population effect of the two common garden locations was estimated together with two variance components, quantifying the site-specific variability in the montane garden and the site-specific variability of the garden location effect (ie lowland – montane). This last term in the model allows us to predict the relative garden performance for each clone compared to the population average. Posterior variance estimates were used to construct 95% confidence intervals for individual random effect predictions. We plotted reaction norms (Schlichting and Pigliucci 1998) of each clone for each of the nine traits to illustrate how the phenotypes of the different genotypes expressed in each of the two gardens.

To visualise the trait differences among the 34 clones and to observe how the same clonal replicates responded in each of the lowland and montane common gardens, we ran a principal component analysis (PCA). Based on the covariance between traits, the scaled and centred clone predictions were summarised in a biplot of the first two principal components. We treated all predictions as new observations, ignoring any further estimation uncertainty. To test for association between all measured traits and elevation we plotted principal component scores vs elevation. We plotted principal component scores vs all environmental variables to test for association between all measured traits and all environmental site environmental variables (Table S1). All statistical analyses were performed using the open-source program R, version 4.0.5 (R Core Team 2018) and package lme4 (Bates et al. 2015).

## RESULTS

### Phenotypic variation

All plants survived in both common gardens and we found considerable variation in all measured traits across the two gardens. In the lowland garden the mean dry weight of the largest clone was almost 20 times that of the smallest, with measures of leaf size varying by more than a twofold difference. As result, there were clear morphotypes; while the majority of plants were somewhat intermediate, individual clones ranged from short, horizontally spreading plants to tall, multi-stemmed and multi flowered individuals. In contrast in the montane garden, vegetative traits among clones were markedly less variable and it was more difficult to recognise morphotypes among the clones. However, in the montane garden flower size varied among clones more than in the lowland garden (Fig. 2). The reaction norms of all clones, except from Park Terrace, Christchurch, responded in a similar way to the environments in the lowland and montane common gardens (Fig. 2).

**Fig 2.**
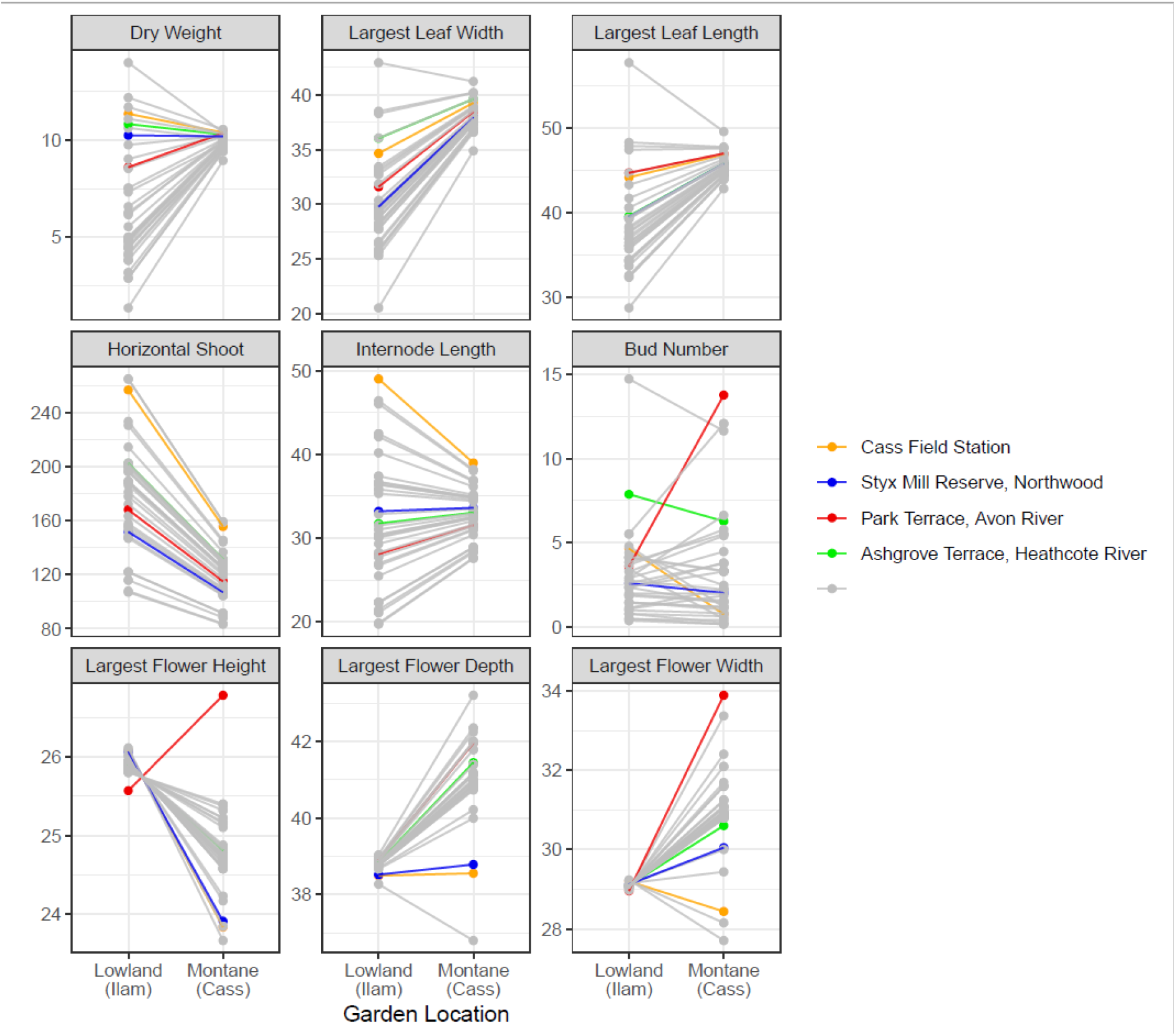
Reaction norms of single genotypes – predictions of expected trait variables by clone. Lines connect the predictions for the same genotype at the two different gardens. All predictions are presented on the original scale of each measurement.

The first two axes of the PCA together explained 66% of the total variation (Fig 3). Axis 1 (38%) correlated most strongly with the following vegetative traits: above ground dry weight, leaf size (length and width), internode length and stolon length. The second axis (28%) correlated most strongly with florescence traits such as flower width and depth. Most variation among clones in the lowland garden was along axis 1, and in the montane garden along axis 2.

**Fig 3.**
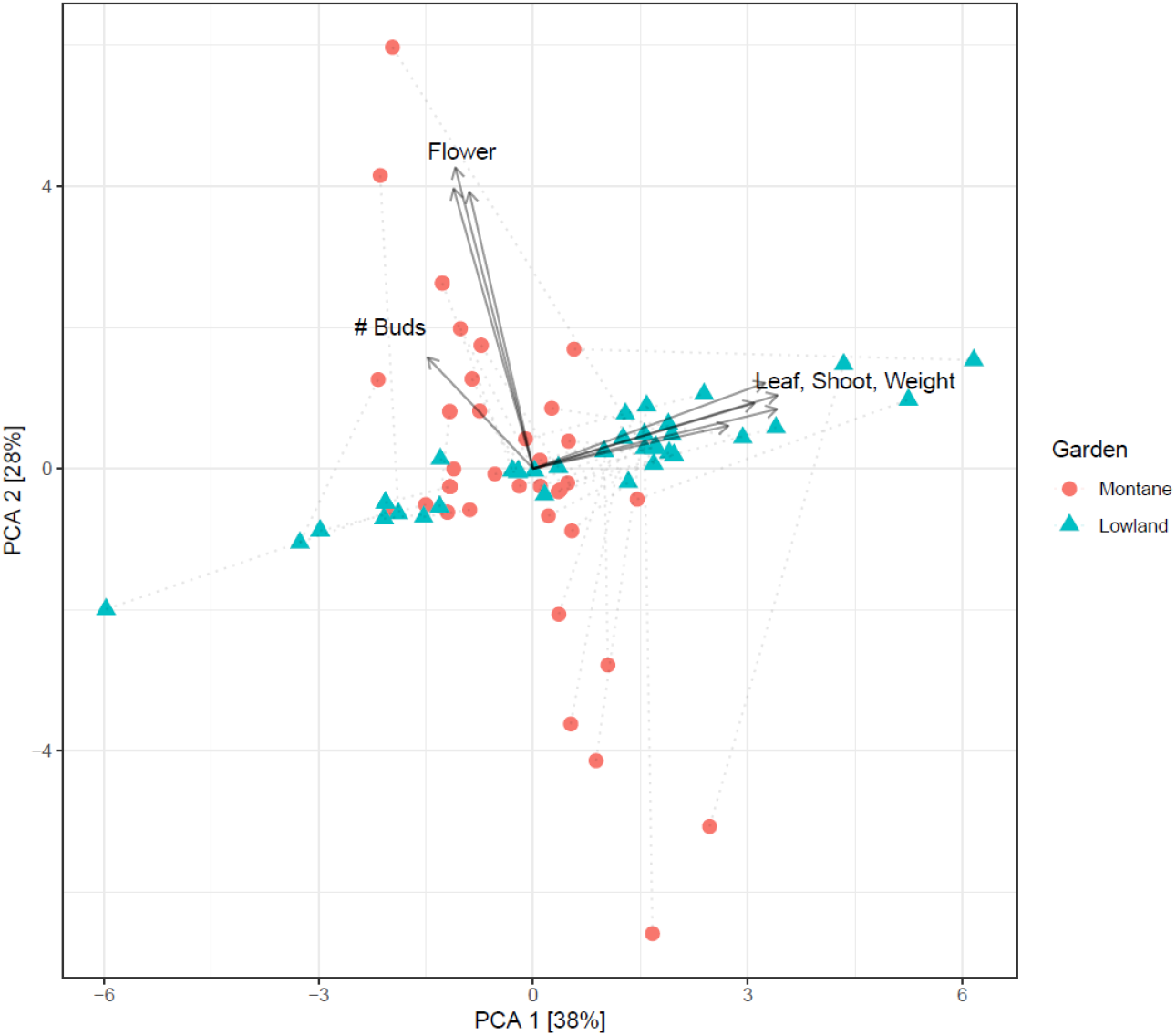
Biplot of the first two principal components, summarising all traits (arrows indicating the rotated coordinate system) for each population-specific prediction. The (0,0) coordinate represents the population average for all traits

All clones show plasticity with variance-changing G x E, and we found marked variation among clones in the degree of plasticity exhibited. For example, two clones from lowland populations in Christchurch showed marked differences in their reaction norms for plant dry weight: the Styx Mill clone showed no shift between elevations while the Park Terrace clone a marked decline in the montane garden (Figure 2). The direction of plasticity also differed among clones, for example despite being collected in the same region and elevation the Styx Mill clone showed a marked decline flower length with elevation while the Park Terrace clone a marked increase in the montane garden. However, while plasticity is common among clones, the rank order of clones remained the same for most traits, except for, the number of buds and flower length where the reaction norm of the Park Terrace was inconsistent with the remaining clones.

Evidence for home site advantage would be apparent if the clone from Cass outperformed all other clones in the montane garden and performed less well relative to lowland clones in the lowland garden. This was generally not the case, within the montane garden the Cass clone is within the top few clones in terms of vegetative traits but is outperformed by several lowland clones. It maintains its relatively high ranking in many traits in the lowland garden. Similarly, there was no evidence at all for the lowland Christchurch sites in the lowland garden outperforming clones from other locales (Fig. 2).

### Genetic versus plastic components to variation

The GLMM analysis to determine total genetic, environmental (residual) and GxE components of the variation in vegetative and reproductive traits among clones revealed that in the lowland garden clone (genotype) explained 33%-45% of the variance in vegetative traits and almost none of the variation in inflorescence traits (Table 2). In contrast in the montane garden it was the inflorescence traits (corolla depth and width and bud number) and stolon length which showed most genetic variability among clones.

**Table 2.** Estimates for site-specific variances by garden location and residual variances together with proportions of explained variance by population [95% confidence intervals]. All estimates are based on logarithmic transformed scales. The total variance among the 34 clones in the Lowland and Montane garden is presented for each measured trait, The residual is the total variance in each measured trait among the 34 clones not explained by garden. The GxE component for each garden is the percentage of variance for each measured trait within each garden

| Trait | $\sigma^2$<br>Montane | $\sigma^2$<br>Lowland | $\sigma^2$<br>Residual | GxE Montane | GxE Lowland |
| --- | --- | --- | --- | --- | --- |
| Dry Weight | 0.01 | 0.80 | 0.99 | 0.01 [0.00; 0.16] | 0.45 [0.30; 0.58] |
| Flower Length | 0.01 | 0.00 | 0.05 | 0.10 [0.00; 0.45] | 0.00 [0.00; 0.29] |
| Flower Depth | 0.01 | 0.00 | 0.02 | 0.21 [0.00; 0.55] | 0.00 [0.00; 0.28] |
| Flower Width | 0.01 | 0.00 | 0.04 | 0.19 [0.00; 0.53] | 0.00 [0.00; 0.28] |
| Leaf Width | 0.00 | 0.06 | 0.08 | 0.04 [0.00; 0.20] | 0.44 [0.28; 0.57] |
| Leaf Length | 0.00 | 0.05 | 0.07 | 0.03 [0.00; 0.15] | 0.41 [0.25; 0.54] |
| Horizontal Shoot | 0.06 | 0.13 | 0.24 | 0.21 [0.06; 0.40] | 0.34 [0.20; 0.48] |
| Internode Length | 0.02 | 0.13 | 0.28 | 0.07 [0.00; 0.23] | 0.33 [0.18; 0.46] |

### Plastic variation

For all of the measured traits except for stolon length and number of buds, the environmental effect of garden was greatest in the montane garden, where almost all of the clones have higher above-ground dry weight, greater leaf areas and longer internode lengths than in the lowland garden (Fig. 4). This environmental component is large, ranging from ~ 20 to 50% difference between gardens, depending on trait.

**Fig 4.**
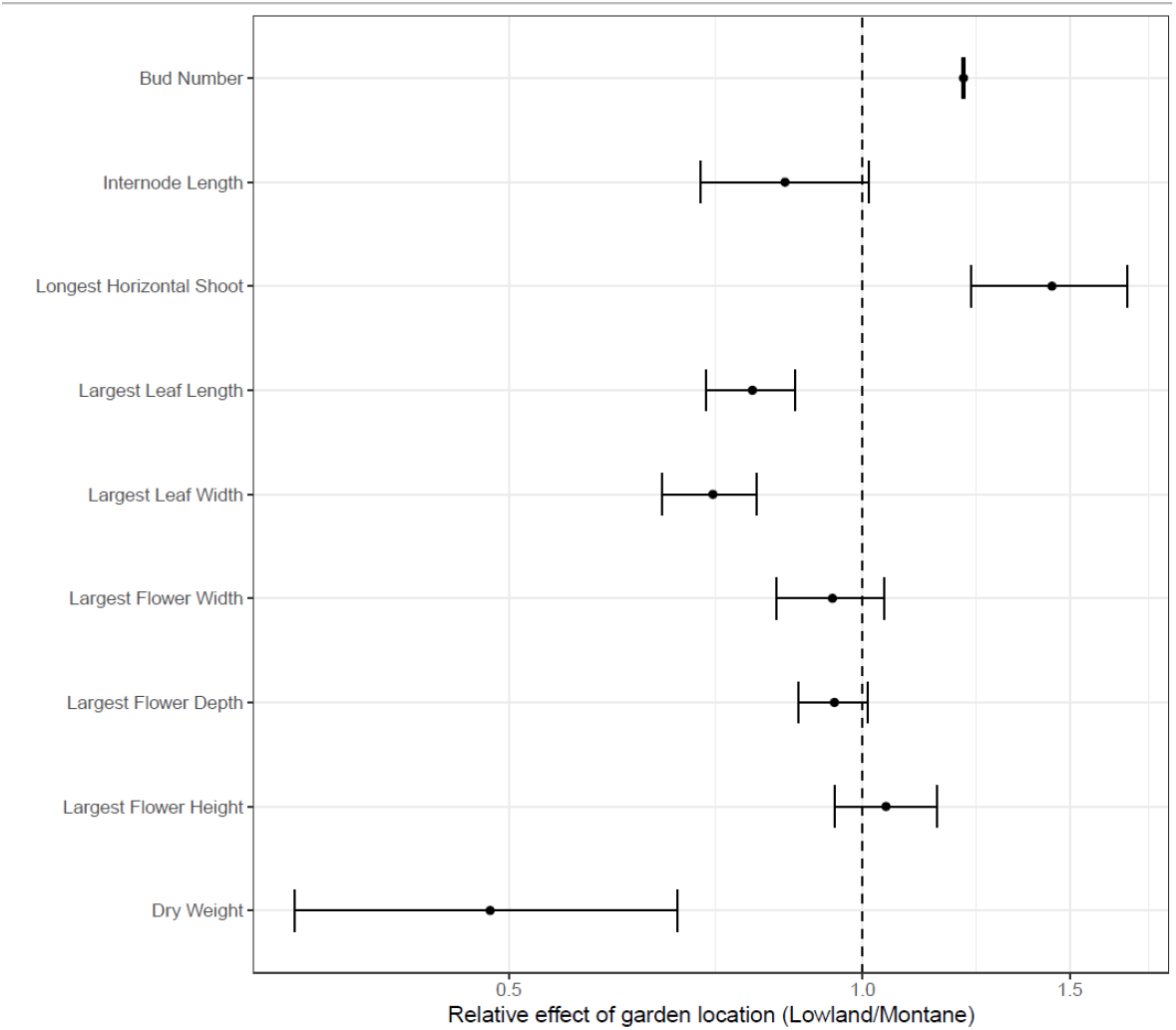
Estimated relative effect of garden location Lowland/Montane and 95% confidence intervals. An estimate of 1 indicates the same expected response at both garden locations, Estimates > 1 correspond to a larger average response in the lowland compared to the montane garden and estimates of <1 indicate a greater average response in the montane, compared to the lowland garden.

The importance of an environmental component to variance is also illustrated by the patterns for a selection of individual clones (Fig. 5, the individual response of each trait in all clones to the two garden environments is presented in Fig 1 Supplementary Information). For example, the clone originating from the montane (Cass) garden site showed one of the strongest and most consistent plastic responses across all traits when moved to the lowland garden, producing longer horizontal shoots and internodes, and more flowers than in its home montane environment (Fig. 5). In contrast, the clones from the lowland Travis Wetland and Ashgrove Terrace sites close to the lowland common garden showed a similar, if less pronounced response to the two garden environments (Fig 5). In the lowland environment stolon length is markedly longer relative to the montane garden in all clones, although the degree of between-garden differences (plasticity) varies among clones. In contrast to the other traits, number of buds are fairly similar in both gardens for most clones, although where there are plastic response these vary in direction among clones. A few clones e.g., Ashgrove Terrace clone, do show a strong environmental response (Fig 5). As expected, we found far more within-clone variation in vegetative and reproductive (length of horizontal shoot and bud number) than floral traits.

**Fig 5.**
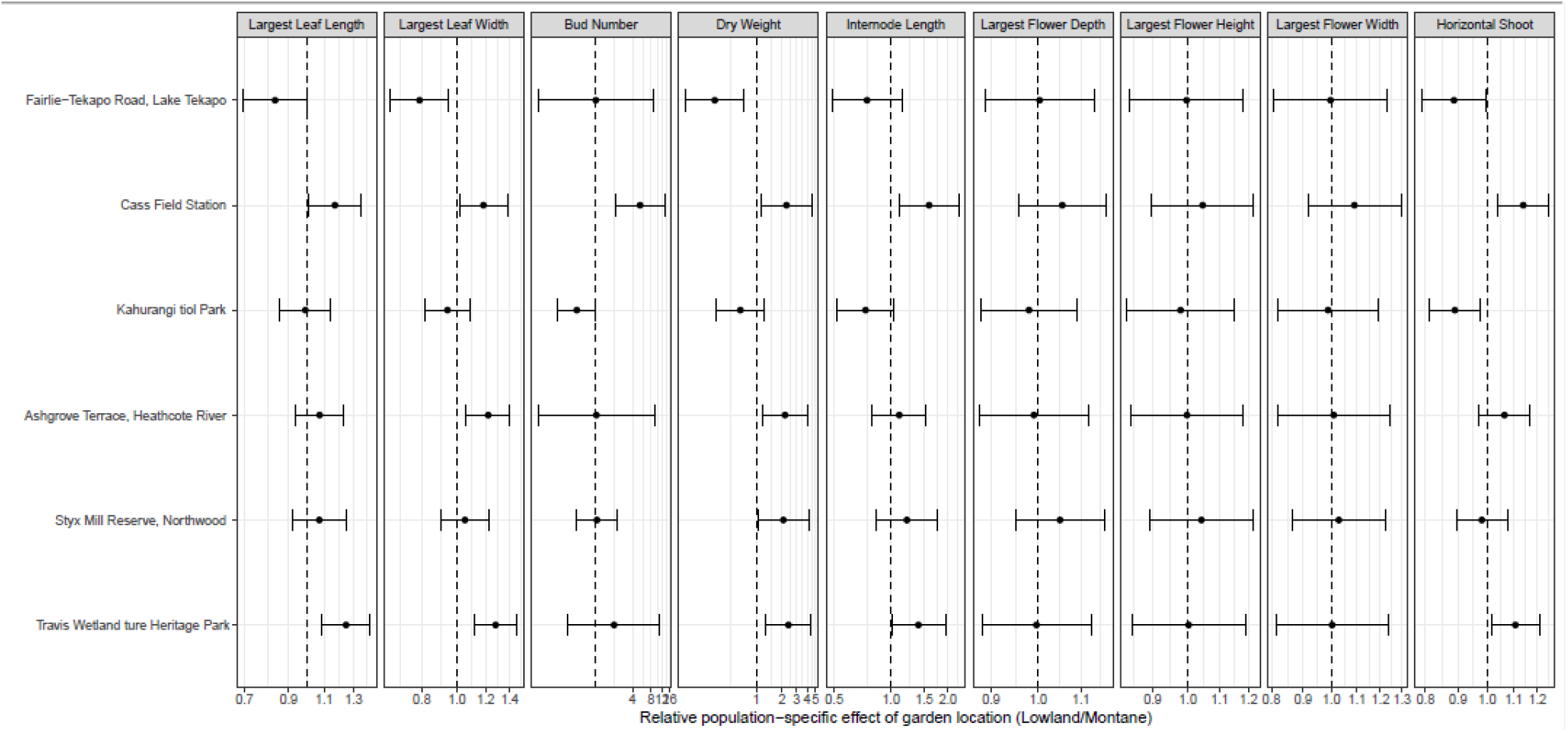
Predicted clone-specific relative effects of garden location Lowland/Montane and 95% confidence intervals. Clones are ordered from highest (990m) to lowest (0m) elevation of the sampling location. A value of 1 indicates the same expected population average at both locations, Estimates > 1 correspond to a larger average response for a clone in the lowland garden compared to montane garden.

### Evidence for adaptation

When the PC scores were plotted against each of the source environmental variables independently, for each clone in each of the lowland and montane garden, elevation explained some of the variation in vegetative traits. This was most obvious in the clones growing in the lowland garden, where the clones from lower elevation source sites were overall markedly larger in comparison to clones from higher elevation source sites (Fig 6).

**Fig 6.**
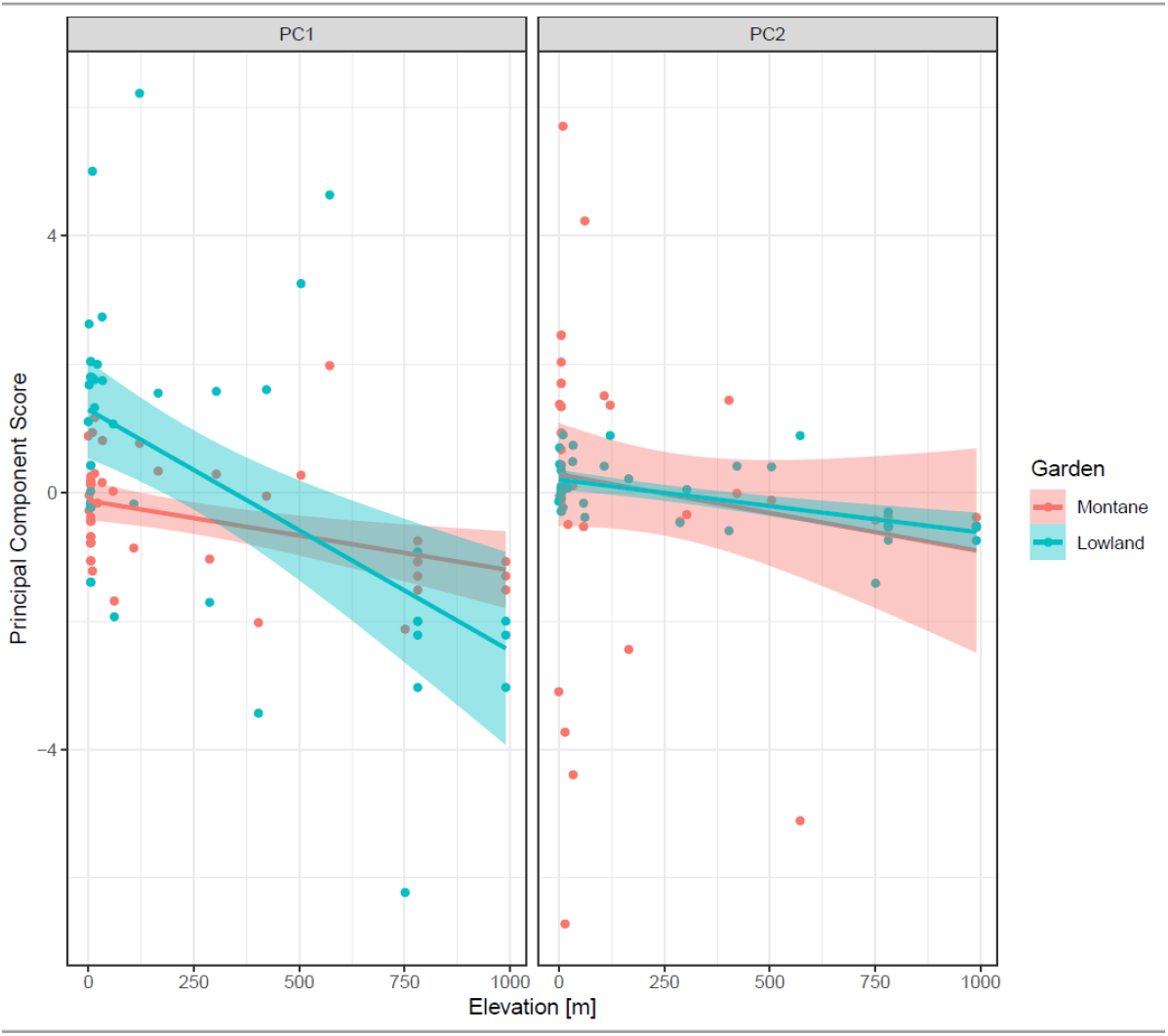
Dependency between Principal Component scores and elevation of sample locations separate for the two garden locations. The predictions of a linear regression model are added.

Clones from lower elevation sites were more plastic than those from higher elevations (Fig. 7). This influence of source elevation on individual clone responses to garden locations is most obvious for above ground dry weight, stolon length and internode length. In contrast, elevation at source location had no marked effect on relative garden response in bud number or flower characteristics (Figure 7).

**Fig 7.**
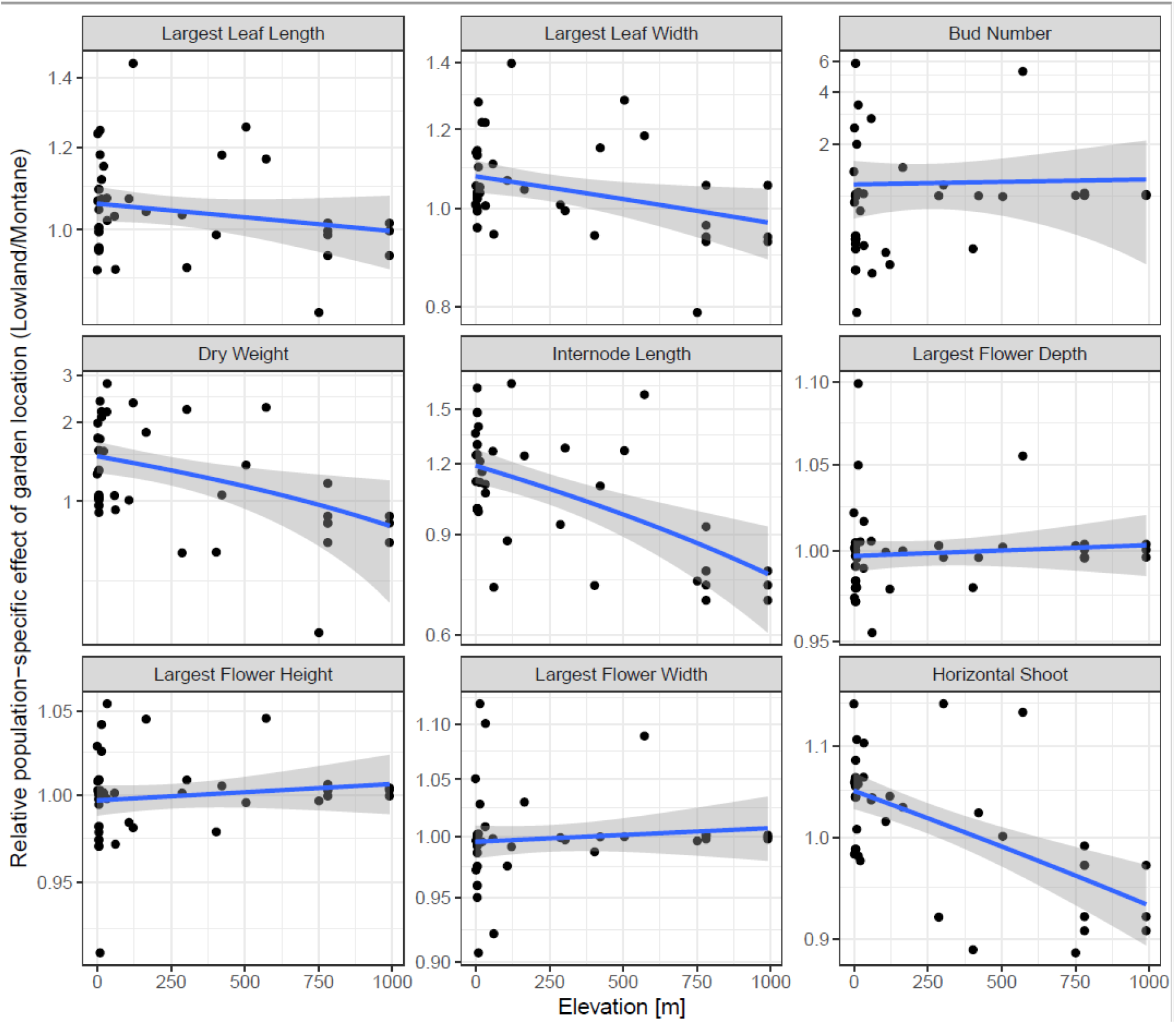
Dependency between the relative clone-specific effect of garden locations and elevation of sample location. The predictions of a (log)-linear model for each of the measured variables is added.

## DISCUSSION

The aim of our study was to describe some of the phenotypic variation in *E. gutatta* clones from across New Zealand and to partition this variation into genetic and plastic components. We hoped to untangle the relative importance of genetic versus plastic effects in the success of *E. gutatta* as an invasive clonal plant, something which might shed light on its modelled niche shift in New Zealand (Da Re et al. 2021). Specifically we tested hypotheses of adaptation to elevation and home-site advantage. While we acknowledge that one clone per site excludes any measure of intra-population variation, we are confident that most sites comprised one or a very few clones.

### Phenotypic variation

Results from our common garden experiment showed that as expected, there were both genetic and environmental components to the variance in all traits measured. However, sorting out the relative contribution of each component and answers about local adaptation became possible only because we used two common gardens and ensured that these gardens were located towards each of the two elevational extremes of source populations (Allan and Pennall 2009; Scheepens et al. 2010; Berend et al. 2019;). We found markedly different results between gardens; in the lowland garden a major proportion of variation in the five measured vegetative traits (above ground dry weight, leaf width, leaf length, horizontal shoot length and internode length) was explained by genetic differences among the clones, while in the montane garden variation in above ground dry weight, leaf width and leaf length was almost entirely masked by plastic responses. In contrast, among-clone genetic differences in flower traits was visible in the montane garden, but hidden in the lowland garden. However, caution is necessary here as conclusions may depend on which fitness measures are used (Villellas et al. 2021). Moreover, plastic responses, some passive and non-adaptive (Liu et al. 2016) especially in vegetative traits, may mask genetic traits under favourable conditions (Villellas et al. 2021).

Previous studies using common garden experiments to explore variation within introduced species in their new range (Williams et al. 2008; Flory et al. 2011), or to investigate home site advantage (Allan and Pennal 2009) have also found marked differences in trait expression among gardens. The more similar the garden environment to the native range, or home site, the better adapted and therefore fitter the plants should theoretically be. We found limited support for a home-site advantage, with plants collected in the environs of each common garden not beings consistently superior to clones from other locations.

### Genotypic variation

In the lowland garden, plants ranged from tall, multiple stemmed, large-leaved individuals with abundant flowers through to low growing clones with long horizontal shoots to small individuals with few flowers. Our model predicts that up to 45% of the variance in plant dry weight and 34 % of the variance in length of horizontal shoots was genetically determined. Taken together, the variance in vegetative, and reproductive traits we observed in the lowland garden suggests that across New Zealand there is extensive genetic variation among *E. gutatta* clones. Results from the montane garden tell a different story; at Cass genetic variation among clones explains only 1% of the variation in dry weight and only 21% of the variation in horizontal shoot length, but more genetic variation in flower traits than was observed in the lowland garden. Basing our understanding on data from the montane garden only would suggest that *E. gutatta* clones across New Zealand show little genetic variation in vegetative traits but some variation in flower traits.

### Fitness differences across environments

Bearing in mind that our proxy for fitness was above ground dry weight, our plots of the PC1 scores against source elevation suggest adaptation to elevation; the plot strongly indicated higher biomass in clones from low elevation compared with high elevation source sites in the lowland garden. Numerous studies globally have demonstrated the adaptive strategy of low plant height and low biomass at high elevations (Körner 2003). Moreover Olsson and Agren (2002) and Monty et al. (2009) have demonstrated this adaptation in introduced plant species in the US and Europe respectively. However, while adaptation is a credible explanation for our finding that clones from lowland sites are fitter in the lowland than in the montane garden, alternative explanations such as genetic drift through founder effects (Monty et al. 2009; McGoey et al. 2020) cannot be dismissed.

The weak signal of advantage in clones from lowland sites in the high elevation Cass garden is in accordance with DeMarche et al. (2016) who found that low elevation *E. gutatta* ecotypes outperformed montane ecotypes in their montane common garden. However, the authors agree their conclusions are tenuous for several reasons, including misappropriate measures of fitness. In our New Zealand study, we did not measure below ground biomass, which may have conferred fitness at higher elevations. Moreover, our results are from one year only so we may have missed inter-annual variation in fitness. Alternatively, it is conceivable that genotypes better adapted to higher elevations in New Zealand have not yet reached high elevations environments.

### Home site advantage

Despite the wide range of genetic variation among our clones in vegetative traits we found little evidence of local adaptation, instead the ranking of clones in terms of relative trait values stayed the same across most traits in both gardens. While the Cass clone, growing in the Cass montane garden was always in the top few ranking clones for all vegetative and reproductive traits we measured, it was only top in internode length and was outperformed by clones from lowland sites in length of longest horizontal shoots-stolons. Similarly, in the Ilam lowland garden while clones from low elevation sites around Christchurch relatively close to the lowland garden performed relatively well they were outperformed by several clones from other regions. Multiple introductions, small populations and founder effects (McGoey et al.), and possibly conflicting selection pressures (Barrett et al. 2008) may explain this lack of evidence for home site advantage. Strong gene flow among populations (Zhao et al. 2013) seems an unlikely explanation given the clonal nature of the spread of *E. gutatta* (Truscott et al. 2008). Multiple introductions into different catchments and founder effects seem the most likely explanation; frequent removal (sometimes annual) of *E. gutatta* to clear waterways followed by reintroduction through fragments of different provenances could well explain lack of ecotypic variation. While absence of ecotypic variation despite strong genotype variation in invasive species has been reported previously (Lord 1992; Ebeling et al. 2011; Pahl et al. 2013; Herden et al. 2019) there is always the possibility that adaptation is being missed because of inappropriate fitness measures. For example, Lord (1992) points out that while Rapson and Wilson (1992) found no evidence for home site advantage among New Zealand *Agrostis capillaris* populations in growth and floral morphology, they did find evidence of local adaption to soil water availability and soil nitrogen and phosphorus. Identifying the key traits is therefore essential (Williams et al. 2008; Bufford and Hulme 2021).

### Plasticity

All clones showed considerable plasticity in all the traits we measured, which was unsurprising, especially for the vegetative traits given this is a perennial, weedy herb (Bazzaz 1996). We found above ground dry weight, our proxy for fitness, was almost universally higher in the montane relative to the lowland garden; dry weight was on average 50% higher across all clones at Cass than at Ilam, with little variance in dry weight among clones at Cass. While this reduced inter-clonal variance in dry weight at montane garden reflected varying steepness of the reaction norm for above ground dry weight among the clones, rank order did not change. Likewise, horizontal shoot length was on average 5% shorter in the montane than in the lowland garden, but with no crossing-over of reaction norms. Consistent reactions norms among clones suggests past adaptation and may be indicative of less potential for local adaptation in new environments than if clones had shown variable reaction norms to different environments (Cheplick 2015). One interpretation of these results therefore is that the montane environment is more similar than lowland Ilam to the environment in the native range of *E. gutatta* populations; plasticity permits clones to maximize their fitness under optimal conditions (Hendry 2016). However, based on the environmental suitability models Da Re et al. (2020), neither the lowland or montane garden are especially suitable areas for *E. gutatta.* It may well be that much of the plasticity we observe across most of the traits we measured is passive, a side effect of adaptations or other plastic responses (De Witt et al. 1998; Funk 2008; Hulme 2008; Auld et al. 2010; Bufford and Hulme 2021a), limiting species ability to respond to changing environments (Valladares et al. 2014). Bufford and Hulme (2021a) stress the need for future experimental studies linking trait plasticity to fitness along manipulated environmental gradients.

The direction of the reaction norm for internode length did vary among clones, with some clones showing longer internode lengths in the montane environment relative to the lowland garden and others shorter. This may indicate adaptive plasticity; certainly foraging behaviour via horizontal and vertical growth as observed in *E. gutatta* should be adaptive by allowing the colonisation of new niches (Clements et al. 2021). van Kleunen and Fischer (2021) tested this hypothesis on the clonally reproducing *Ranunculus reptans* and found genetic variation in plasticity for internode and stolon length, evidence supporting adaptive plasticity. The importance in our study of these results is that they show genetic variation among clones for a plastic trait which may support invasiveness (Clements et al. 2021).

Further evidence for adaptive (active) plasticity was found in bud number. For this trait reaction norm slopes crossed in some of the clones, and different clones responded differently to the two gardens. Again, this illustrates the potential for adaptive evolution across our *E. gutatta* clones.

In contrast to the other traits, the flower length, depth and width showed considerable among clone genetic variation in the montane environment but elicited a similar plastic response to the lowland garden, where flowers from all clones were essentially similar in size. We caution too much interpretation of these results because of low flower numbers, but plasticity of *E. gutatta* flowers has been reported before, a response to drought (Kelly et al. 2008). That clones from lower elevation source sites showed greater plasticity in vegetative traits than clones from high elevation source sites is important. Plasticity is selected for in changing environments (Oostra et al. 2018) and lowland sites may be more varied environments than montane ones, experiencing more fluctuation in nutrient levels and more anthropogenic disturbance. Whatever the reason for this elevational variation the finding is notable because the distribution of plastic traits across species ranges may impact how species respond to new environments and how they affect range shifts (Bufford and Hulme 2021a).

### Future work

Our study illustrates the potential opportunity in using *E. gutatta* to further understand the mechanisms of successful plant invasions into new niches. Key to taking this further is experimentation using multiple clones from each population in reciprocal transplants, both within New Zealand and between New Zealand and native US *E. gutatta* populations. Comparing home and away fitness traits and home and away plasticity (Bufford and Hulme 2021b), especially in traits associated with clonal spread will be important. Trade-offs (or not) associated with greater horizontal shoot length and *e.g.* flower production and seed set should be explored. To what extent plasticity may be aiding or confounding future adaptation and spread across New Zealand needs further investigation. Future fitness trait measures should include flowering phenology, well known to influence invasion success in plants and physiological traits which may be more closely associated with fitness than the traits included in this study.

### Conclusions

Our study has shown the presence of considerable genetic variation among *E. gutatta* clones from across the South Island of New Zealand; while we found scant evidence for local adaptation (home site advantage) we found a strong signal for higher fitness in all vegetative traits across both garden environments in clones from lowland, as compared with higher elevation sites. Clones from lowland sites were more plastic relative to clones from higher elevation sites, including two key traits facilitating the spread of clonal plants, horizontal shoot length and internode length. Having variation for plasticity in essential traits such as these suggests that *E. gutatta* is well adapted to move into new environments (Gratani 2014) and that plasticity may contribute substantially to its successful niche expansion within New Zealand as compared with its native range. To what extent this reflects a trade-off or lack thereof with other reproductive traits (Bufford and Hulme 2021b) is a major focus for future investigation.

## Acknowledgments

The study was funded through grants from the Koiata Botanical Trust with MW receiving a Roland Stead Postgraduate Scholarship and a Laura J Clad Memorial Scholarship.

Many thanks to Dave Conder for maintaining the Ilam garden and to Aaron Millar for field work support.

## Data Availability Statement

All data are available on request

## Statements & Declarations

### Author Contributions

All authors contributed to the study conception and design. Material preparation, data collection was mainly by M.W. with contributions from H.C. Data analysis were performed by Daniel Gerhard. The main draft of the manuscript was written by Hazel Chapman and all authors commented on previous versions of the manuscript. All authors read and approved the final manuscript.

