## Supplemental Table 1 for "Relative roles of genetic variation and phenotypic plasticity in the invasion of monkeyflower *Erythranthre gutatta* in New Zealand"

*Table S1: The environmental variables recorded or measured for each of the 35 populations across New Zealand. Temperatures were measured in "℃", altitude measured in "m", rainfall and precipitation in "mm". *0 means no, 1 means yes. **Environmental averages for the growing period (September – March) 1971-2000*

| Region | Location Name | Population | Latitude | Longitude | Water Logged* | Flowing Water* | Altitude (m) | Annual Average Temperature | Average Temperature** | Average Min Temperature** | Extreme Min Temperature** | Average Max Temperature** | Extreme Max Temperature** | Annual Average Rainfall | Humidity** | Precipitation** | Temperature at Collection | Sunlight Hours |
|---|---|---|---|---|---|---|---|---|---|---|---|---|---|---|---|---|---|---|
| SI NW | Kawatiri-Murchison Highway | 1 | -41.695 | 172.478 | 0 | 0 | 271 | 12.36 | 15.16 | 8.26 | -1.55 | 22.08 | 32.85 | 871.6 | 74.4 | 70.6 | 25 | 15.03 |
| SI NW | St Arnaud-Kawatiri Highway | 2 | -41.697 | 172.637 | 0 | 1 | 271 | 9.88 | 12.48 | 6.13 | -5.7 | 18.83 | 30 | 871.6 | 74.4 | 70.6 | 24 | 15.03 |
| SI NE | Taylor River, Blenheim 1 | 3 | -41.509 | 173.954 | 1 | 1 | 7.55 | 13.33 | 15.53 | 10.26 | 0.17 | 20.79 | 31.64 | 433.9 | 78.7 | 30.9 | 28 | 15.01 |
| SI NE | Hawkesbury Road, Hawkesbury | 4 | -41.524 | 173.817 | 1 | 1 | 59.25 | 12.68 | 14.98 | 8.88 | -1.94 | 21.04 | 32.69 | 433.9 | 78.7 | 30.9 | 28 | 15.01 |
| SI NE | Taylor River, Blenheim 2 | 5 | -41.512 | 173.960 | 1 | 1 | 8.6 | 13.33 | 15.53 | 10.26 | 0.17 | 20.79 | 31.64 | 433.9 | 78.7 | 30.9 | 28 | 15.01 |
| SI NE | Waikawa Road, Waikawa | 6 | -41.273 | 174.033 | 1 | 1 | 18.56 | 13.54 | 15.51 | 11.64 | 2.3 | 19.4 | 27.61 | 433.9 | 78.7 | 30.9 | 28 | 15.01 |
| SI NE | Rapaura Road, Spring Creek | 7 | -41.459 | 173.959 | 1 | 1 | 8.23 | 13.33 | 15.53 | 10.26 | 0.17 | 20.79 | 31.64 | 433.9 | 78.7 | 30.9 | 17 | 15.01 |
| SI NE | Hillocks Road, Spring Creek | 8 | -41.457 | 173.950 | 1 | 1 | 8.77 | 13.33 | 15.53 | 10.26 | 0.17 | 20.79 | 31.64 | 433.9 | 78.7 | 30.9 | 17 | 15.01 |
| SI NW | Kahurangi National Park | 9 | -41.685 | 172.441 | 0 | 1 | 250.63 | 12.36 | 15.16 | 8.26 | -1.55 | 22.08 | 32.85 | 871.6 | 74.4 | 70.6 | 24 | 15.01 |
| SI NW | Upper Buller Gorge | 10 | -41.811 | 172.064 | 0 | 1 | 204.4 | 12.36 | 15.16 | 8.26 | -1.55 | 22.08 | 32.85 | 871.6 | 74.4 | 70.6 | 24 | 15.01 |
| SI C | Fairlie-Tekapo Road, Lake Tekapo | 11 | -44.007 | 170.488 | 1 | 1 | 714.63 | 9.01 | 12.01 | 3.09 | -4.7 | 18.53 | 30.86 | 493.1 | 76.9 | 40.5 | 26 | 15.25 |
| SI C | Tekapo-Twizel Road, Tekapo | 12 | -44.175 | 170.323 | 1 | 1 | 519.71 | 9.82 | 13.24 | 5.35 | -5.6 | 21.13 | 33.15 | 493.1 | 76.9 | 40.5 | 26 | 15.24 |
| SI C | Tekapo Twizel Track, Pukaki | 13 | -44.254 | 170.116 | 1 | 1 | 457.22 | 9.82 | 13.24 | 5.35 | -5.6 | 21.13 | 33.15 | 493.1 | 76.9 | 40.5 | 26 | 15.24 |
| SI C | Omarama-Lindis Pass Road, Waitaki 6 | 14 | -44.505 | 169.781 | 1 | 1 | 576.41 | 9.82 | 13.05 | 6.1 | -4.2 | 19.99 | 33.03 | 493.1 | 76.9 | 40.5 | 26 | 15.27 |
| SI C | Omarama-Linids Pass Road, Waitaki 2 | 15 | -44.532 | 169.712 | 1 | 1 | 664.48 | 9.82 | 13.05 | 6.1 | -4.2 | 19.99 | 33.03 | 493.1 | 76.9 | 40.5 | 26 | 15.27 |

| | | | | | | | | | | | | | | | | | | |
|---|---|---|---|---|---|---|---|---|---|---|---|---|---|---|---|---|---|---|
| SI C | Omarama-Lindis Pass Road, Waitaki 10 | 16 | -44.565 | 169.662 | 1 | 1 | 664.48 | 9.82 | 13.05 | 6.1 | -4.2 | 19.99 | 33.03 | 493.1 | 76.9 | 40.5 | 26 | 15.27 |
| SI C | Omarama-Lindis Pass Road, Waitaki 4 | 17 | -44.506 | 169.782 | 1 | 1 | 664.48 | 9.82 | 13.05 | 6.1 | -4.2 | 19.99 | 33.03 | 493.1 | 76.9 | 40.5 | 26 | 15.27 |
| SI C | Fairlie-Tekapo Road, Mackenzie | 18 | -44.066 | 170.672 | 0 | 0 | 501.14 | 10.6 | 12.71 | 6.82 | -1.98 | 17.04 | 31.03 | 493.1 | 76.9 | 40.5 | 26 | 15.25 |
| SI C | Geraldine-Fairlie Highway | 19 | -44.097 | 170.834 | 1 | 0 | 298.31 | 10.6 | 12.71 | 6.82 | -1.98 | 17.04 | 31.03 | 493.1 | 76.9 | 40.5 | 26 | 15.25 |
| SI SE | Waihola Highway, Milburn | 21 | -46.074 | 170.013 | 1 | 1 | 44.53 | 10.37 | 12.69 | 6.57 | -3.56 | 18.3 | 31.81 | 1080.5 | 79.2 | 92.9 | 16 | 15.43 |
| SI SE | Owaka Highway, Clutha | 22 | -46.377 | 169.690 | 1 | 0 | 60 | 9.99 | 12.03 | 7.02 | -2.08 | 17.03 | 28.53 | 1080.5 | 79.2 | 92.9 | 12 | 15.47 |
| SI SE | Owaka Highway, Katea, Otago | 23 | -46.419 | 169.693 | 0 | 0 | 99.03 | 10.33 | 11.95 | 8.33 | 0.78 | 15.58 | 27.56 | 1080.5 | 79.2 | 92.9 | 12 | 15.47 |
| SI SE | Papatowai Highway, Owaka | 24 | -46.459 | 169.646 | 0 | 0 | 75 | 10.33 | 11.95 | 8.33 | 0.78 | 15.58 | 27.56 | 1080.5 | 79.2 | 92.9 | 16 | 15.47 |
| SI SE | Papatowai Highway, Clutha | 25 | -46.487 | 169.545 | 1 | 0 | 177.44 | 10.35 | 11.96 | 7.39 | -0.13 | 16.56 | 29.04 | 1080.5 | 79.2 | 92.9 | 16 | 15.47 |
| SI SE | Tokanui-Gorge Road Highway, Fortrose | 26 | -46.560 | 168.790 | 0 | 0 | 4.26 | 10.77 | 12.41 | 8.78 | 0.24 | 15.39 | 19.85 | 1080.5 | 79.2 | 92.9 | 26 | 14.46 |
| SI SW | Te Anau-Milford Highway, Southland | 27 | -45.137 | 167.931 | 0 | 1 | 339.92 | 9.29 | 11.66 | 5.25 | -3.26 | 16.47 | 28.32 | 644.8 | 66.3 | 54.2 | 23 | 15.38 |
| SI CE | Styx Mill Reserve, Northwood | 29 | -43.464 | 172.609 | 1 | 1 | 11.92 | 11.64 | 14.12 | 8.71 | -2.66 | 19.53 | 33.12 | 131.3 | 75.3 | 10.1 | 25 | 15.25 |
| SI CE | Keating Street, Silverstream | 30 | -43.379 | 172.634 | 1 | 1 | 6.44 | 11.67 | 13.92 | 8.4 | -1.22 | 19.44 | 32.7 | 131.3 | 75.3 | 10.1 | 25 | 15.25 |
| SI CE | Jeffs Drain Road, Ohoka | 31 | -43.394 | 172.598 | 1 | 1 | 10.14 | 11.67 | 13.92 | 8.4 | -1.22 | 19.44 | 32.7 | 131.3 | 75.3 | 10.1 | 25 | 15.25 |
| SI CE | Travis Wetlands Heritage Park | 32 | -43.485 | 172.699 | 0 | 1 | 2.05 | 12.29 | 14.7 | 9.63 | -0.62 | 19.76 | 32.5 | 131.3 | 75.3 | 10.1 | 25 | 15.25 |
| SI CE | Park Terrace, Avon River | 33 | -43.523 | 172.627 | 1 | 1 | 9.69 | 12.29 | 14.7 | 9.63 | -0.62 | 19.76 | 32.5 | 131.3 | 75.3 | 10.1 | 25 | 15.25 |
| SI CE | Ashgrove Terrace, Heathcote River | 34 | -43.567 | 172.627 | 1 | 1 | 10.22 | 12.29 | 14.7 | 9.63 | -0.62 | 19.76 | 32.5 | 131.3 | 75.3 | 10.1 | 25 | 15.25 |
| SI C | Cass Field Station | 36 | -43.035 | 171.758 | 1 | 1 | 578.81 | 8.22 | 10.72 | 4.17 | -5.3 | 17.27 | 30.06 | 2289.6 | 84 | 192.6 | 19 | 14.45 |
| SI SW | Fiordland National Park, Fiordland | 37 | -45.101 | 167.968 | 0 | 1 | 279.77 | 8.22 | 10.72 | 5.25 | -3.26 | 16.47 | 28.32 | 644.8 | 66.3 | 54.2 | 23 | 15.38 |
